# Checklist of the birds of Wanang Conservation Area, Papua New Guinea

**DOI:** 10.1101/2025.08.21.671428

**Authors:** Katerina Sam, Krystof Korejs, Samuel Jeppy, Bonny Koane

**Affiliations:** Biology Centre of the Czech Academy of Sciences, Institute of Entomology, Branisovska 31, Ceske Budejovice 370 05, Czech Republic; Faculty of Science, University of South Bohemia, Branisovska 1760, Ceske Budejovice 370 05, Czech Republic; The New Guinea Binatang Research Centre, Madang, Papua New Guinea

## Abstract

The Wanang Conservation Area in Papua New Guinea represents a large, community operated protected lowland rainforests. Based on surveys conducted in both the wet (November) and dry (July) seasons between 2010 and 2011, we report a total of 147 bird species known in the area around Swire station. Based on these surveys, we provide abundances of 120 bird species recorded during the standardized point-counts and mist-netting. Further, we add notes on the observations of another 27 bird species which we detected in the region outside of the standardized surveys. Finally, we compare our results with previous (2009) and later shorter surveys conducted in the region (2013 and 2019). In total, we document a cumulative total of 153 bird species recorded in the area to date.

## Introduction

New Guinea, the world’s largest tropical island (>750,000 km^2^), is renowned for its exceptional avian diversity and remarkable number of endemic bird lineages (Pratt and Beehler 2015, Beehler and Pratt 2016). The island harbours the third-largest contiguous primary rainforest globally (Brooks et al. 2010) and supports over five percent of the world’s forest biodiversity (Sekhran and Miller 1995). More than 800 bird species inhabit New Guinea, with species richness peaking at low elevations (Pratt and Beehler 2015, Beehler and Pratt 2016, Sam et al. 2019).

Until the early 20th century, New Guinea’s lowland rainforests were largely spared from large-scale commercial logging (Fox et al. 2011). In Papua New Guinea (PNG), which occupies the island’s eastern half, over 70% of primary forests have retained their integrity to that time (Gamoga et al. 2021), providing extensive undisturbed habitats for birds and other taxa. Yet, in recent decades, deforestation and forest degradation have accelerated sharply (Bryan et al. 2015, Turia et al. 2022), particularly in lowland regions where biodiversity is most at risk (Shearman and Bryan 2011). Despite the growing threats, scientific knowledge about the impacts of habitat loss on PNG’s rich avifauna remains limited. The establishment of large, formal conservation areas has been slow, and much of the ongoing conservation effort is driven by initiatives led by local communities.

The Wanang Conservation Area (WCA, > 10,000 ha), situated in the heart of the lowland tropical rainforest of Madang Province, Papua New Guinea, exemplifies community-based conservation efforts in the region (Novotny and Toko 2015). Managed locally by a community of about 200 inhabitants, the WCA protects one of the most intact tracts of lowland rainforest in the province. In addition to its importance for ecosystem services and carbon storage, the area has become a hub for ecological research. Nevertheless, biodiversity assessments remain fragmentary and unsystematic, despite numerous intensive studies conducted over the past two decades (e.g., Klimes et al. 2011, Vincent et al. 2015, Chmel et al. 2016, Chmel et al. 2017, Vinagre□Izquierdo et al. 2022, Ezedin 2023, Korejs et al. 2023).

Ornithological surveys of varying duration and methodological rigor have also been carried out at WCA during the last 15 years (Tvardíková 2010, Chmel et al. 2016, Chmel et al. 2017, Hazell et al. 2021), yet a complete checklist of the area’s avifauna has not been published. Regular field courses in tropical ecology, jointly organized by the University of South Bohemia (Czech Republic) and the New Guinea Binatang Research Center (Papua New Guinea) every couple of years, further contribute to research and training in ornithology. During these courses, students, assisted by local guides, often survey bird communities, and their results have been published, incrementally enriching knowledge of the local bird assemblages (Tvardíková 2010, Korejs et al. 2023).

This checklist seeks to bridge the current gap in systematic documentation of bird records from the Wanang Conservation Area (WCA), focusing on intense surveys conducted between 2010 and 2011 and placing these observations in the context of earlier and later published studies. Our data from point-count bird surveys and mist-netting document the abundances of individual bird species and highlight several species of conservation concern. The objectives of this checklist are: 1) To provide a comprehensive taxonomic inventory of bird species encountered in WCA, and 2) to establish a baseline for future monitoring, ecological research, and conservation management decisions by the Wanang community. By compiling and presenting these records, we aim to advance knowledge of Papua New Guinea’s avifauna and to support long-term conservation in one of the island’s last extensive tracts of intact lowland rainforest.

## Methods

### Study sites

We conducted the field work in Wanang Conservation Area in Papua New Guinea in the proximity of the Wanang 3 (Swire station) research station (5.2277608S, 145.0797250E) and its 50 ha Forest Dynamic Plot (FDP, which is located centrally within WCA). The site is located within the continuous primary forest, in the middle of Wanang Conservation Area, which is a ∼10,000 ha area covered by largely continuous, lowland rainforest, which is selectively logged at patches close to borders. The climate is characterized by a mean annual temperature of 25.8 °C and annual precipitation of 4000 mm, has a mild dry season from July to September. (McAlpine et al. 1983, Vincent et al. 2015). The detailed data presented here are based on the surveys we conducted in November 2010, July 2011 and November/December 2011, and the list obtained is compared by species lists of previously published works (Chmel et al. 2016, Hazell et al. 2021, Korejs et al. 2023).

### Bird surveys

We surveyed the bird communities using point counts and mist-netting. The counts were conducted at 16 points semi-regularly (ca. 150-200 m apart) spaced along a 2250-m transect (Figure 1, Table 1). We started censuses 15 min before sunrise (ca. 5:45 AM) at a randomly selected starting point, and we then continued counts in a randomly selected direction. We counted birds for 15 minutes at each point, so all 16 points were surveyed before 11:00 AM. We consider one such census on all 16 points to be one replication in time.

**Table 1.**
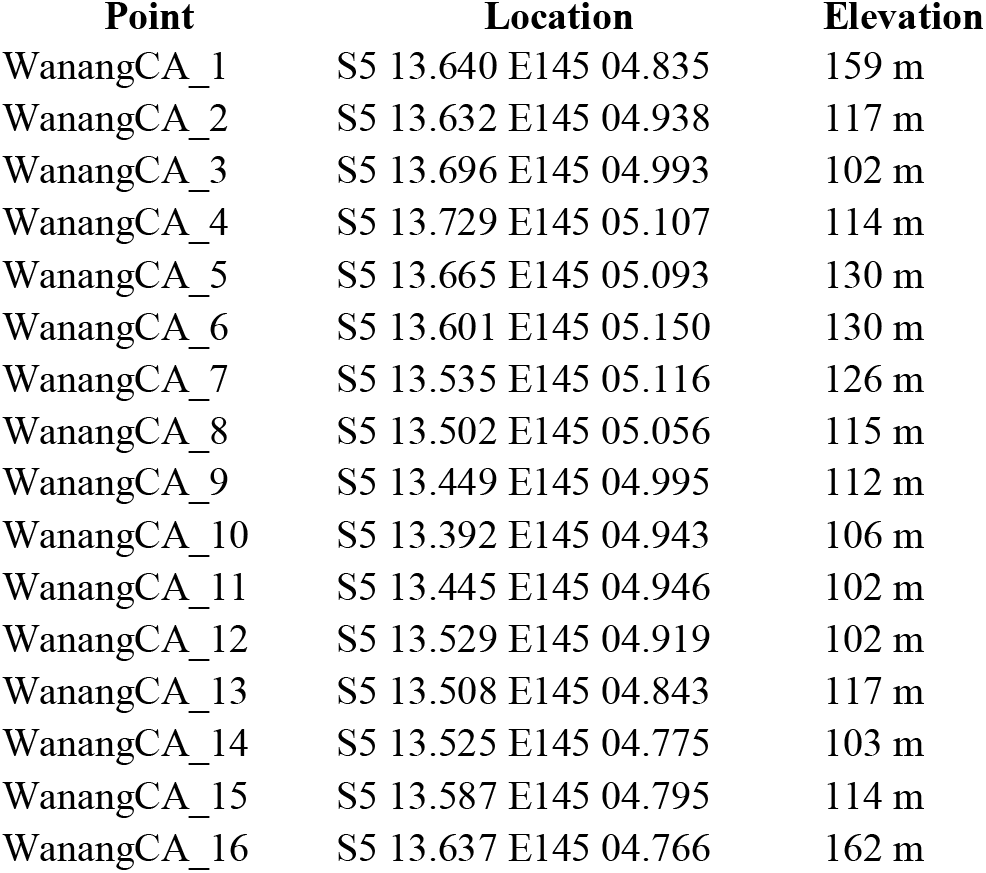
Locations of the 16 points on which the census was conducted during point-counts in 2010-2011.

| Point | Location | Elevation |
| --- | --- | --- |
| WanangCA_1 | S5 13.640 E145 04.835 | 159 m |
| WanangCA_2 | S5 13.632 E145 04.938 | 117 m |
| WanangCA_3 | S5 13.696 E145 04.993 | 102 m |
| WanangCA_4 | S5 13.729 E145 05.107 | 114 m |
| WanangCA_5 | S5 13.665 E145 05.093 | 130 m |
| WanangCA_6 | S5 13.601 E145 05.150 | 130 m |
| WanangCA_7 | S5 13.535 E145 05.116 | 126 m |
| WanangCA_8 | S5 13.502 E145 05.056 | 115 m |
| WanangCA_9 | S5 13.449 E145 04.995 | 112 m |
| WanangCA_10 | S5 13.392 E145 04.943 | 106 m |
| WanangCA_11 | S5 13.445 E145 04.946 | 102 m |
| WanangCA_12 | S5 13.529 E145 04.919 | 102 m |
| WanangCA_13 | S5 13.508 E145 04.843 | 117 m |
| WanangCA_14 | S5 13.525 E145 04.775 | 103 m |
| WanangCA_15 | S5 13.587 E145 04.795 | 114 m |
| WanangCA_16 | S5 13.637 E145 04.766 | 162 m |

**Figure 1.**
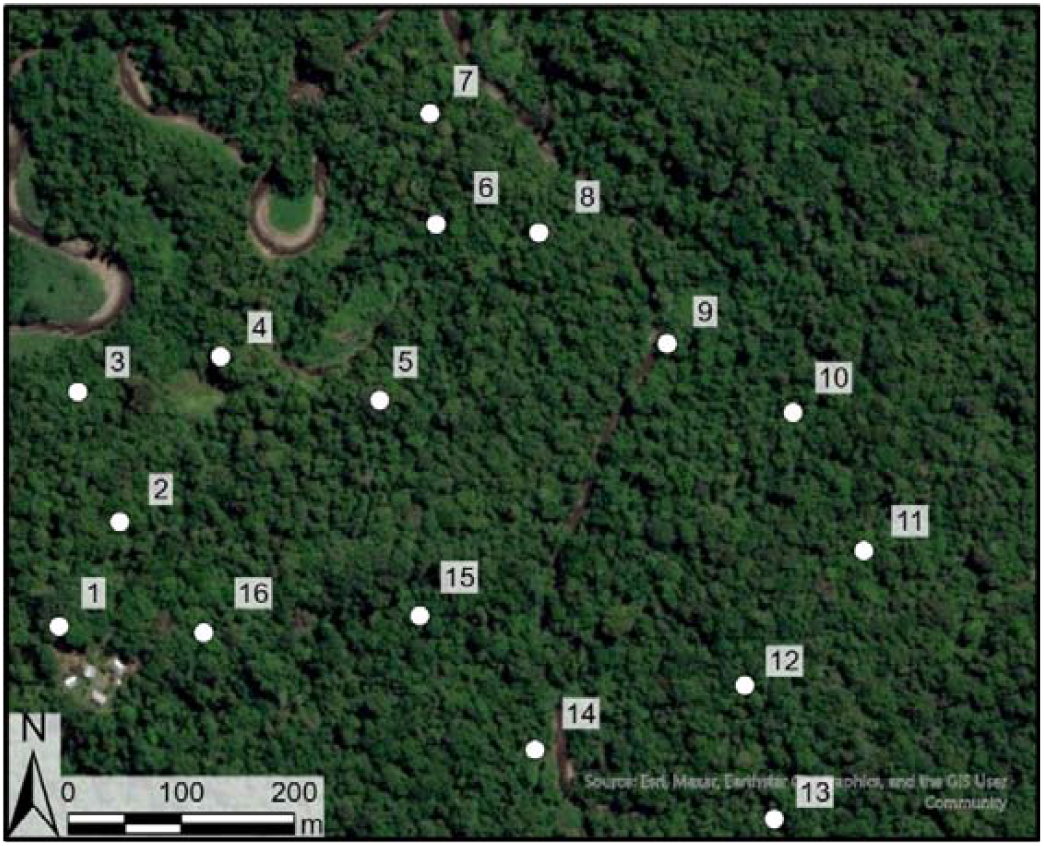
Map showing the location of the 16 points in the proximity of the Swire research station (i.e., so called Wanang 3) in the Wanang Conservation Area. Exact GPS locations of the points are in Table 1.

During the surveys, we recorded all birds seen or heard within 50 m radius of the point (more detailed surveys were done and used for other purposes than this species list, see e.g. for methodology (Sam et al. 2019). To minimize double-counting, we attempted to accurately track movements of birds and recorded more individuals of the same species only when they called at the same time or from distinctively different directions. In November 2010 we conducted 9 replicates of the censuses in time (4^th^ till 17^th^), in July 2011 again 9 replicates (between 5^th^ and 21^st^) and in November-December 2011 we conducted again 9 replicates (between 24^th^ and 3^rd^) in time. All surveys were conducted by KS, BK, and SJ, all of whom had previous experience with bird surveys in Papua New Guinea.

In addition to point-counts, we conducted mist-netting in three blocks of 6 days during each of the visits we made to Wanang. We mist-netted birds along a 200-m-long line of nets (2.5 m high × 12– 18 m long each, 16-mm mesh) for 12 hours daily from 05:30 AM to 17:30 PM, with checks every 20 minutes. During the first 3 days, nets were placed between the first three points of the point-count transect, and then transferred to the last three points of the point-count transect for the next 3 d. We identified all mist-netted birds to species, marked them with colour bands, and released them within 10 minutes. On top of these standardized surveys, we recorded bird species continuously during our stays, during our other activities – i.e., during cooking, walking, washing in river. These additional bird species are recorded without densities.

## Results

All together, we recorded 8,851 individuals of birds during point-counts and 230 birds during mistnetting in Wanang Conservation Area during the standardized surveys in 2010 and 2011. These birds represented 120 species (Table 2). On top of these bird species recorded during the standardized times, we also recorded additional 27 bird species outside of these surveys, in our free time. Altogether, we thus report 147 bird species from the Wanang Conservation Area, from the proximity of the Wanang 3 research station based on this survey. The bird species observed outside of our standardized surveys are following species:

**Table 2.**
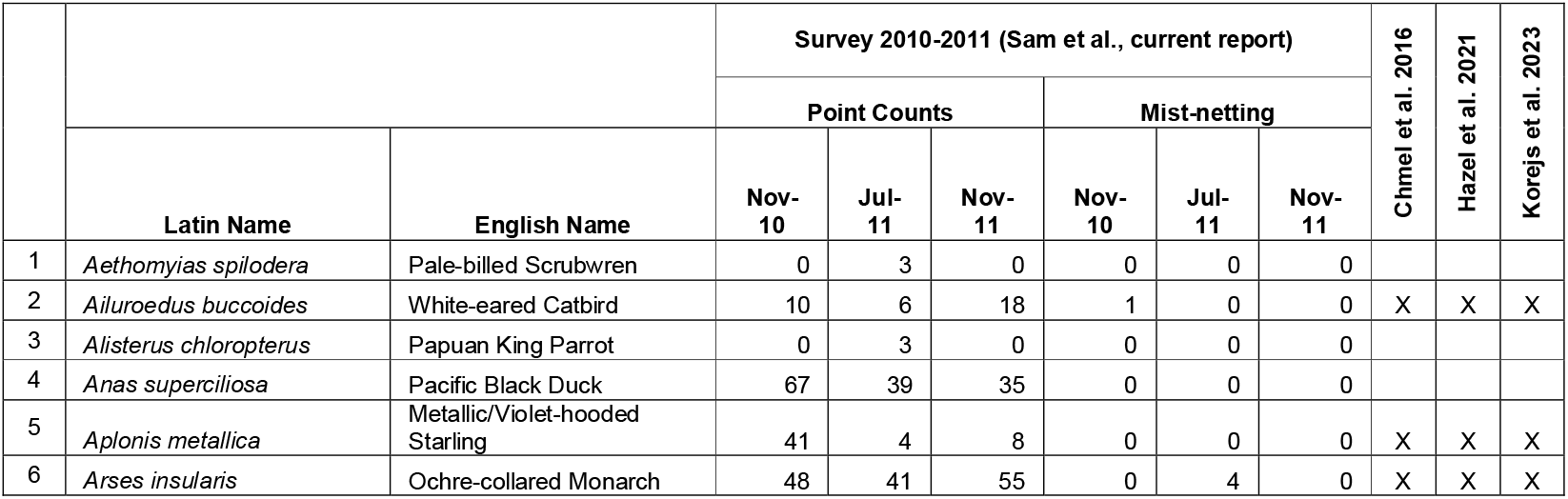
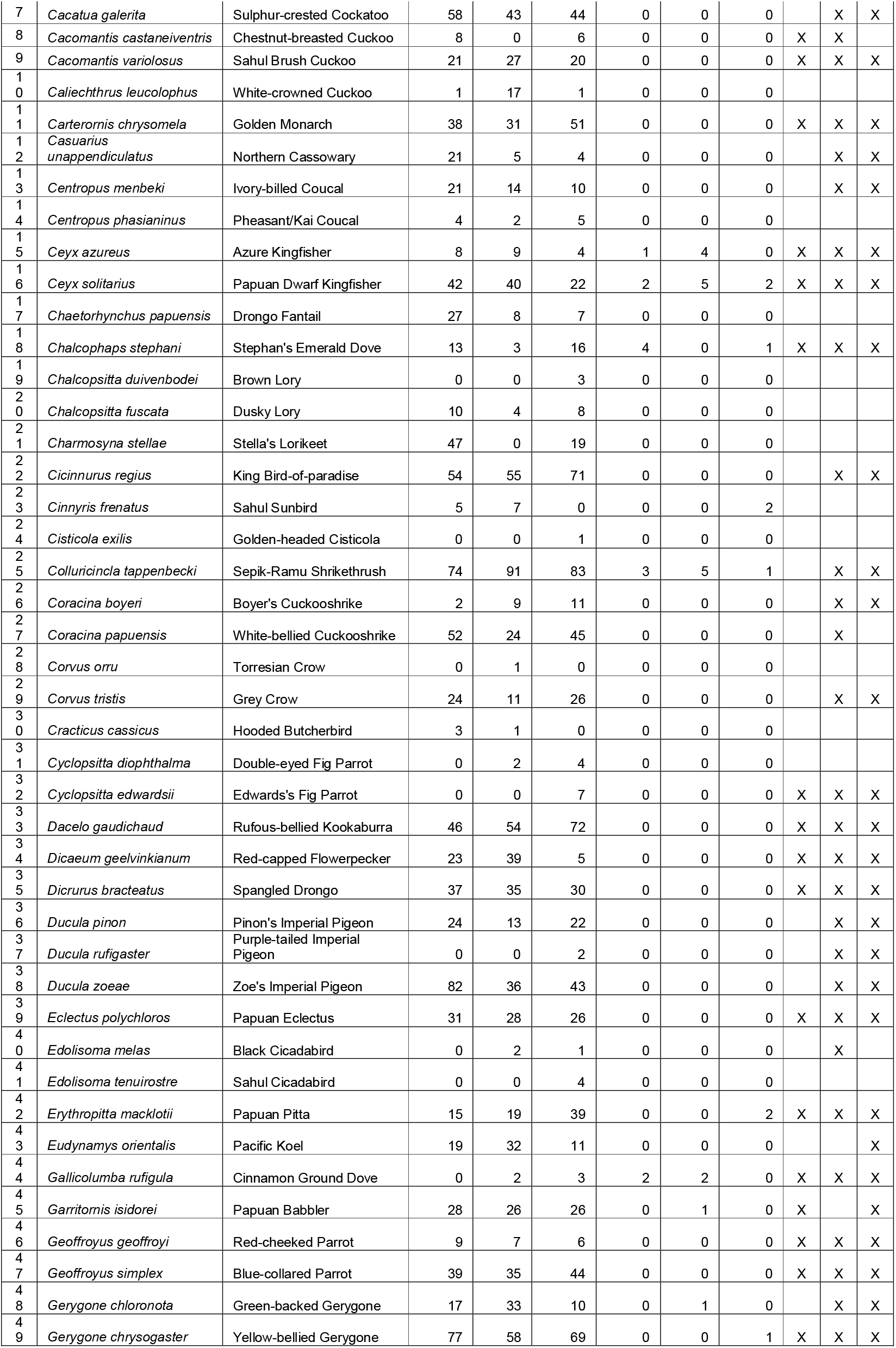
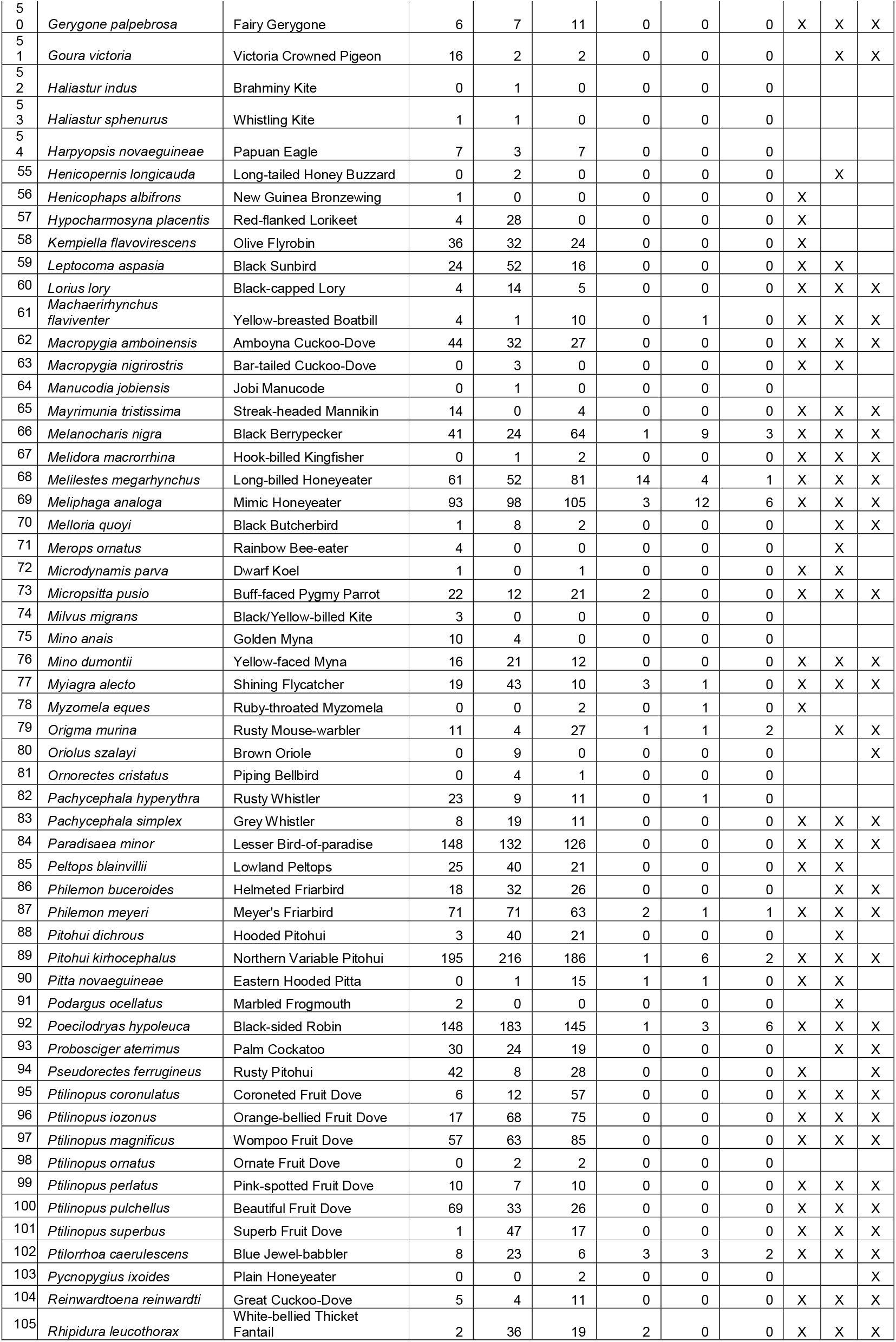
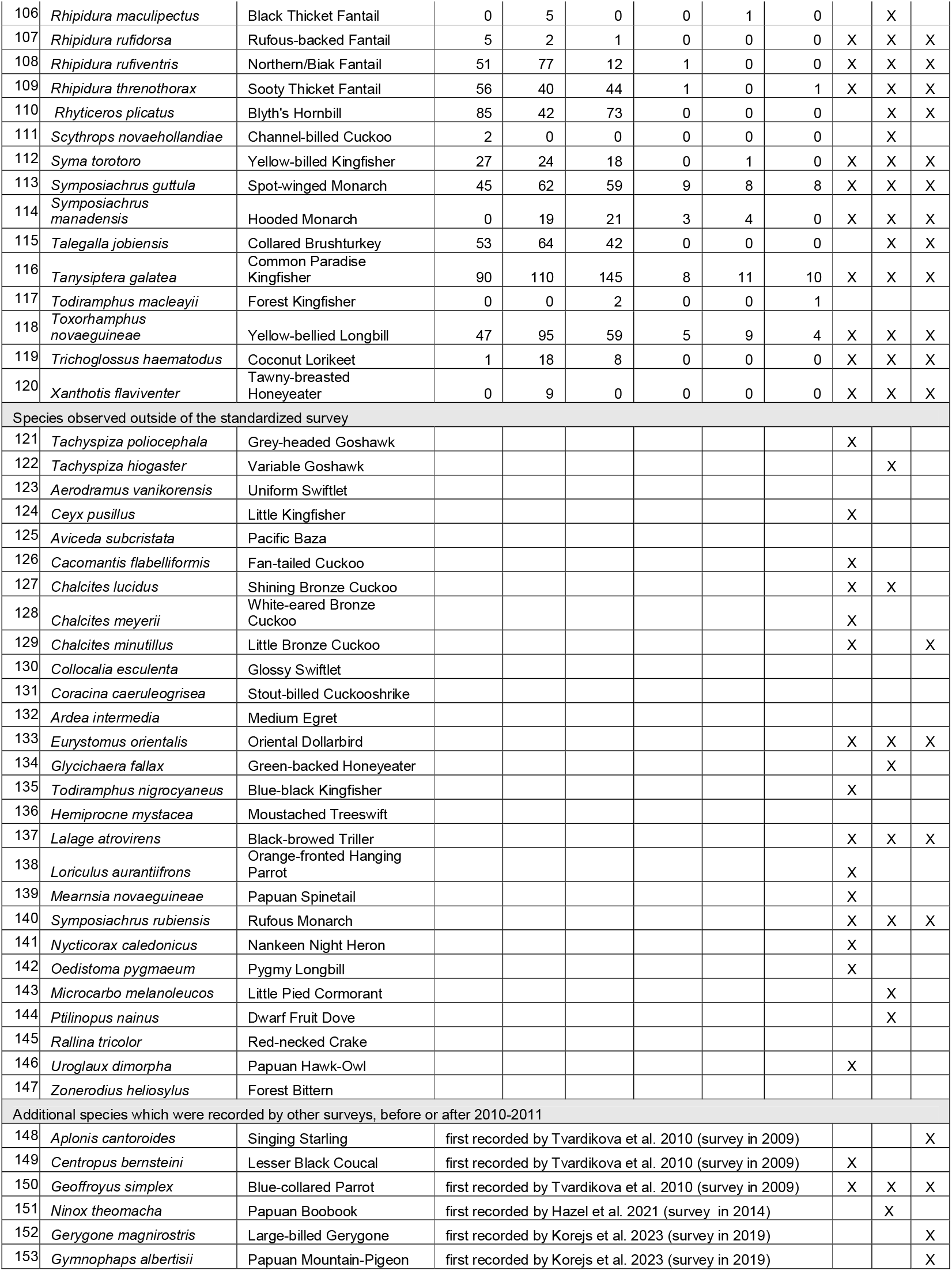
List of the bird species and their abundances recorded during the 27 replications of point-counts (3x 9 days on 16 points) and during 18 days of mist-netting conducted in 2010-2011. Additionally, presences of the species in other published results of the surveys are marked by X, in the last three columns. Within the table, all birds except two have LC (Least concern status). Two exceptions are underlined and marked as VU – Vulnerable or NT – Near threatened.

Grey-headed Goshawk (*Tachyspiza poliocephala*,, LC) – Observed once, in the forest gap in the 50ha Dynamic Forest Plot. Reported also by Chmel et al. 2016.

Variable Goshawk (*Tachyspiza hiogaster*, LC) – observed as one fly over above the area, in July 2010.

Uniform Swiftlet (*Aerodramus vanikorensis*, LC) – commonly flying over, during all stays, typically not recorded during point-counts due to canopy coverage.

Little Kingfisher (*Ceyx pusillus*, LC) – observed three times during the July 2010 survey along the riverbanks.

Pacific Baza (*Aviceda subcristata*, LC) – observed as flying above in November 2011.

Fan-tailed Cuckoo (*Cacomantis flabelliformis*, LC) – observed once in July 2010 and once in July 2011. Reported also by Chmel et al. 2016.

Golden Cuckooshrike (*Campochaera sloetii*, LC) – observed up to 6 individuals, in November 2010, in the afternoon in the proximity of the points 1 and 16, i.e., in the forest gap around the station.

Shining Bronze Cuckoo (*Chalcites lucidus*, LC) – observed twice in July 2011.

White-eared Bronze Cuckoo (*Chalcites meyerii*, LC) – observed several times in July 2011, flying around the riverbanks. Reported also by Chmel et al. 2016.

Little Bronze Cuckoo (*Chalcites minutillus*, LC) – observed several times in July 2010 and 2011.

Reported also by Chmel et al. 2016, and by Korejs et al. 2023

Glossy Swiftlet (*Collocalia esculenta*, LC) – common fly-over the camp, hard to see through forest canopy.

Stout-billed Cuckooshrike (*Coracina caeruleogrisea*, LC) – observed once in November 2011, potentially heard in the following days by the local assistants.

Medium Egret (*Ardea intermedia*, LC) – seen often at the riverbanks.

Oriental Dollarbird (*Eurystomus orientalis*, LC) – observed twice in the helipad forest gap in July and November 2011.

Green-backed Honeyeater (*Glycichaera fallax*, LC) – heard and seen repeatedly in the afternoons in July 2010, along the riverbank of the helipad. In Papua New Guinea, we were more often recording the species in forest fragments and in secondary forests, so this seems to be a rare observation in the primary forest. Reported also by Chmel et al. 2016.

Blue-black Kingfisher (*Todiramphus nigrocyaneus*, NT) – observed in July 2010 along the access road to the research station from the Wanang 2 river. Reported also by Chmel et al. 2016.

Moustached Treeswift (*Hemiprocne mystacea*, LC) – common fly-over the study site.

Black-browed Triller (*Lalage atrovirens*, LC) – a female observed over the three consecutive days in November 2011 along the riverbank. Reported also by Chmel et al. 2016.

Orange-fronted Hanging Parrot (*Loriculus aurantiifrons*, LC) – observed four times, three times in November 2010 and once in November 2011. Reported also by Chmel et al. 2016.

Papuan Spinetail (*Mearnsia novaeguineae*, LC) – observed several times flying over the canopy. Reported also by Chmel et al. 2016.

Rufous Monarch (*Symposiachrus rubiensis*, LC) – observed three times in Wanang site, on 2^nd^ and two individuals on 6^th^ of December 2010 (in the proximity of points 11 and 14, in the afternoon).

Nankeen Night Heron (*Nycticorax caledonicus*, LC) - observed once in the area, flying along the river.

Pygmy Longbill (*Oedistoma pygmaeum*, LC) – observed commonly in the afternoons in the forest gap of the helipad and the station.

Little Pied Cormorant (*Microcarbo melanoleucos*, LC) – observed repeatedly flying along the river.

Dwarf Fruit Dove (*Ptilinopus nainus*, LC) – observed once, on the access road to the camp in November 2011.

Red-necked Crake (*Rallina tricolor*, LC) – observed walking along the helipad at the dusk in July 2011.

Papuan Hawk-Owl (*Uroglaux dimorpha*, LC) – heard repeatedly at hights during all surveys. Reported also by Chmel et al. 2016.

Forest Bittern (*Zonerodius heliosylus*, NT) – on served several times in the area of the 50ha Forest Dynamic Plot.

## Discussion

This checklist (Table 2) represents result of the several robust bird surveys conducted in Wanang Conservation Area in Papua New Guinea between the years 2010 and 2011. Note that much larger area of WCA is yet to be surveyed. Despite its local importance, the species list will potentially aid further research in the area. The results could assist the survey of the changes in bird communities and potential future responses of birds to the changes in forest usage or climatic change, as the forest and area surveyed is likely representative to the rest of the conservation area. Despite considerable survey effort, a number of additional species might be still expected to occur in the area and have to be still recorded. As a student course of Tropical Ecology takes place in the WCA regularly, some of the student surveys indeed revealed new bird records (e.g., Table 2 – Korejs et al. 2023), contributing further to the knowledge on the avifauna of the area.

As an example of such student work during the student course, the birds were surveyed at similar locations as reported here for the first time in 2009. This short survey reported 91 bird species. Out of these 91 previously recorded bird species, four species were not confirmed by our research conducted in 2020 and 2011. The unconfirmed species were Singing Starling (*Aplonis cantoroides*) which is a species typical for more open areas, and has a distinctive black colour, unlike the shining metallic colour of Metallic/Violet-hooded Starling (*Aplonis metallica*). Other unconfirmed species was Black-billed Coucal (*Centropus bernsteini*) which is generally known from the region but easily missed species. Two other Centropus species recorded in 2010-2011 have very distinctive sounds, and these three species are very difficult to misidentified when heard (which was the case for all records). Another species which was observed in the survey in 2009 and not confirmed in 2010-2011 was a highly mobile, Blue-collared Parrot (*Geoffroyus simplex*). Finally, previously reported Varied Triller (*Lalage leucomela*) was not confirmed in 2010-2011 surveys, and might have been actually mistaken with Black-browed Triller (*Lalage atrovirens*). Despite the calls of the species are distinct, female individuals were observed in all instances, and they are rather similar, unlike the male individuals. In general, our robust surveys conducted in 2010-2011 confirmed 87 out of 91 bird species reported in 2009 and added additional 62 bird species. Three species (but not *Lalage atrovirens*) were however recorded by the later surveys (Table 1) in the region, which only shows that even intense surveys might miss some rare and illusive species.

Another study, focusing on the vertical stratification of the Wanang forest species was conducted in 2013 by Chmel and his team (Chmel et al. 2016), and reported 84 bird species. All 84 bird species described by this study have been previously recorded in the region by our survey in 2010-2011. With respect to problematic Lalage species, the study of Chmel et al. 2016 reports only *Lalage atrovirens* but not *Lalage leucomela*, so it is possible that original identification was indeed wrong.

The study by Korejs et al. (2023), conducted in July/August 2019, further extended the list of the species know in WCA. The study has shown records of Papuan Mountain Pigeon (*Gymnophaps albertisii*), an otherwise montane species of pigeon which reportedly exhibits seasonal migration to lowland rainforests. In addition, the work reports Large-billed Gerygone (*Gerygone magnirostris*) which shows that even lowland species continue to be found within the region, despite no previous detection from 2009-2011. The presence of the Large-billed Gerygone in the WCA is confirmed by a mist-netting effort conducted by a Danish team working in Swire station in 2016, when the team captured a single specimen. This specimen is located now in the National History Museum (catalogue number NHMD217608).

This would mean that in the primary forests of the Wanang Conservation Area, 147 bird species were recorded in years 2010 and 2011, with 3 more species recorded in 2009 (Tvardikova et al. 2010), and additional 1 in 2014 (Hazel et al. 2021) and 2 more species in 2019 (Korejs et al. 2023), totalling thus to 153 bird species. We predict that further surveys might still reveal more species at the site, as we might be missing mostly nocturnal birds, raptors and birds which prefer to stay in the natural gaps within the primary forests. For examples, the assistants report an observation of Long-billed Cuckoo (*Chalcites megarhynchus*) which was reportedly observed by visiting scientists form Australia (pers. com B. Koane) and unconfimed records of Black-fronted/Green-fronted White-eye (*Zosterops minor*), which might be a seasonal visitor. Overall, our report contributes to the knowledge of the birds of Wanang Conservation Area, which is still far from complete.

## Acknowledgements

We are grateful to Vojtěch Novotný for introducing us to the Wanang Conservation Area through the tropical ecology courses organized by the University of South Bohemia. Our deepest thanks go to the Wanang community for granting us permission to conduct research on their land and for welcoming us over more than a decade of repeated visits. We also thank the local assistants whose invaluable help made the ornithological surveys possible. The compilation of this species list was supported by various funding sources, primarily the Czech Science Foundation, and was prepared specifically with support provided to KS and KK under grant 22-17593M.

## References

Beehler, B. M., and T. K. Pratt. 2016. Birds of New Guinea: Distribution, Taxonomy, and Systematics. Princeton University Press.

Brooks, T. M., R. A. Mittermeier, G. A. da Fonseca, J. F. Lamoreux, C. G. Mittermeier, and J. Gerlach. 2010. Global biodiversity conservation priorities: an expanded review. A handbook of environmental management. Edward Elgar Publishing.

Bryan, J., P. Shearman, G. Aoro, F. Wavine, and J. Zerry. 2015. The Current State of PNG’s Forests and Changes Between 2002 & 2014.

Chmel, K., J. Riegert, L. Paul, P. Fibich, J. Leps, K. Molem, G. Weiblen, and V. Novotny. 2017. Fine-scale spatial patterns in a tropical forest bird community: topography matters. University of South Bohemia, Ceske Budejovice.

Chmel, K., J. Riegert, L. Paul, and V. Novotný. 2016. Vertical stratification of an avian community in New Guinean tropical rainforest. Population Ecology 58:535–547.

Ezedin, Z. 2023. Contributions to floristics in New Guinea and species delimitation in the Wanang Forest Dynamics Plot. Unpublished PhD thesis, University of Minnesota–Twin Cities, St. Paul, USA.

Fox, J. C., R. Keenan, C. Brack, and S. Saulei. 2011. Native forest management in Papua New Guinea: advances in assessment, modelling and decision-making.

Gamoga, G., R. Turia, H. Abe, M. Haraguchi, and O. Iuda. 2021. The forest extent in 2015 and the drivers of forest change between 2000 and 2015 in Papua New Guinea: deforestation and forest degradation in Papua New Guinea. Case Studies in the Environment 5:1442018.

Hazell, R. J., K. Chmel, J. Riegert, L. Paul, B. Isua, G. S. Kaina, P. Fibich, K. Molem, A. J. Stewart, and M. R. Peck. 2021. Spatial scaling of plant and bird diversity from 50 to 10,000 ha in a lowland tropical rainforest. Oecologia 196:101–113.

Klimes, P., M. Janda, S. Ibalim, J. Kua, and V. Novotny. 2011. Experimental suppression of ants foraging on rainforest vegetation in New Guinea: testing methods for a whole□forest manipulation of insect communities. Ecological Entomology 36:94–103.

Korejs, K., J. Riegert, M. Kigl, and V. Novotny. 2023. Differences in bird community structure between riparian and upland zones in a New Guinean rainforest. Australian Field Ornithology 40:179–195.

McAlpine, J. R., R. Keig, and R. Falls. 1983. Climate of Papua New Guinea. CSIRO and Australian National University Press, Canberra.

Novotny, V., and P. Toko. 2015. Ecological research in Papua New Guinean rainforests: insects, plants and people. The State of the Forests of Papua New Guinea 2015. Measuring change over period 2002-2014:71-85.

Pratt, T. K., and B. M. Beehler. 2015. Birds of New Guinea. Princeton University Press.

Sam, K., B. Koane, D. C. Bardos, S. Jeppy, and V. Novotny. 2019. Species richness of birds along a complete rain forest elevational gradient in the tropics: Habitat complexity and food resources matter. Journal of Biogeography.

Sekhran, N., and S. Miller. 1995. Papua New Guinea country study on biological diversity. Department of Environment and Conservation, Conservation Resource Centre.

Shearman, P., and J. Bryan. 2011. A bioregional analysis of the distribution of rainforest cover, deforestation and degradation in Papua New Guinea. Austral Ecology 36:9–24.

Turia, R., G. Gamoga, H. Abe, V. Novotny, F. Attorre, and L. Vesa. 2022. Monitoring the multiple functions of tropical rainforest on a national scale: an overview from Papua New Guinea. Case Studies in the Environment 6:1547792.

Tvardíková, K. 2010. Bird abundances in primary and secondary growths in Papua New Guinea: a preliminary assessment. Tropical Conservation Science 3:373–388.

Vinagre□Izquierdo, C., K. H. Bodawatta, K. Chmel, J. Renelies□Hamilton, L. Paul, P. Munclinger, M. Poulsen, and K. A. Jønsson. 2022. The drivers of avian□haemosporidian prevalence in tropical lowland forests of New Guinea in three dimensions. Ecology and evolution 12:e8497.

Vincent, J. B., B. Henning, S. Saulei, G. Sosanika, and G. D. Weiblen. 2015. Forest carbon in lowland P apua N ew G uinea: Local variation and the importance of small trees. Austral Ecology 40:151–159.

